# Molecular mechanism of water and glycerol transport through hydrophobic selectivity filter in the aquaporin homolog of *Trypanosoma brucei*

**DOI:** 10.64898/2026.04.27.720980

**Authors:** Peerzada Maajid Parsa, Ramasubbu Sankararamakrishnan

## Abstract

The protozoan parasite *Trypanosoma brucei* is implicated in deadly African sleeping sickness. Experimental studies show that *T. brucei* codes for three aquaporin homologs (TbAQP1 to TbAQP3). TbAQP2 has been established as the high affinity drug transporter of drugs pentamidine and melarsoprol. Mutation in TbAQP2 or its loss result in pentamidine-melarsoprol cross-resistance. TbAQP2 is also shown to transport water, glycerol and other solutes to respond to osmoregulation in the infected hosts or glycerol metabolism. Experimentally determined structures of TbAQP2 shows that it adopts the same aquaporin-like hourglass helical fold. However, the so called aromatic/arginine selectivity filter (Ar/R SF) in TbAQP2 has neither arginine nor aromatic residue and all four residues are hydrophobic. Mutation and functional studies have demonstrated the role of Ar/R SF residues in the transport and selectivity of solutes in aquaporin homologs. The intriguing question is how the completely hydrophobic Ar/R SF region enables the transport of water and glycerol molecules. In this study, we used computational approach to elucidate the molecular mechanism of water and glycerol transport. Our equilibrium molecular dynamics simulations showed that the number of water molecules transported by TbAQP2 is almost one order of magnitude higher than that of prototype water channel AQP1. Moreover, the residence time within TbAQP2 channel is much less compared to that found in AQP1. The relatively wider constriction, interactions of water molecules with the selectivity filter residues and the contact duration, all contribute to a large number of water molecules transported through TbAQP2 channel. Our umbrella sampling studies show that when glycerol is transported through TbAQP2, it participates in interactions with channel residues that can be considered as complimentary to that observed in prototype glycerol transporter GlpF. Our studies reveal the molecular mechanism of water and glycerol transport in TbAQP2 and establish that TbAQP2 is an efficient water transporter.

**Statement of Significance:** *Trypanosoma brucei* causes African sleeping sickness and a homolog of aquaporin, TbAQP2, is involved in the transport of drugs that are used to treat this disease. Developing anti-parasitic drugs requires the knowledge of molecular mechanism of the protein’s function. TbAQP2 has been shown to transport water and glycerol. Permeating solutes have to pass through a narrow constriction region formed by all hydrophobic residues. In the present study, equilibrium molecular dynamics simulations showed that TbAQP2 transports water molecules faster in large quantity in comparison with mammalian AQP1. Higher water transport is due to relatively wider constriction and minimum water interactions with selectivity filter hydrophobic residues. Permeating glycerol molecule is involved in complementary interactions with the channel residues. Our studies reveal how water and glycerol are transported through hydrophobic selectivity filter in TbAQP2.

## Introduction

Members of Major Intrinsic Protein (MIP) superfamily include aquaporins and aquaglyceroporins and are present from bacteria to humans (1–2). MIP homologs are also identified in many different types of parasites including those that cause serious diseases in humans (3–4). They play diverse roles in affecting the physiology of the infected organisms. These MIP homologs are attractive targets for developing anti-parasitic drugs. *Trypanosoma brucei* is one such parasite which is known to cause African sleeping sickness (5). *T. brucei* has three aquaporin homologs (TbAQP1 to TbAQP3) which are closely related to mammalian AQP3 and AQP9 with 40 – 45% sequence identity (6). Expression of TbAQPs in *Xenopus oocytes* exhibited increased permeability of water, glycerol and dihydroxyacetone indicating that the role of these channels in osmoregulation and glycerol metabolism in *T. brucei* (7–8). Melarsoprol and pentamidine are among the few drugs that are used to treat African sleeping sickness in humans (9–10) and the cross-resistance to these drugs are linked to loss of function of TbAQP2 (11). There are conflicting studies that have indicated either TbAQP2 to be a high affinity transporter of pentamidine or the drugs inhibit the glycerol permeability of TbAQP2 (12–13). Recently, two independent studies reported the structure of TbAQP2 using cryo electron microscopy technique and the structures are almost identical. Chen et al. showed that the drugs pentamidine and melarsoprol bound to TbAQP2 channel and suggested that the drugs permeate via TbAQP2 channel (14). TbAQP2 structures determined by Matusevicius et al. showed the drugs and glycerol bound inside the channel (15). Structures of aquaporins and aquaglyceroporins from MIP superfamily have been determined from diverse organisms including *E.coli*, yeast, plants and mammals (1). Although they are sequentially diverse, all MIP structures show a characteristic hourglass helical fold with six transmembrane segments (TM1 to TM6) and two half-helices (LB and LE) formed by the loops connecting TM2-TM3 and TM5-TM6. Examination of channel structures indicates that there are two constrictions: one is called aromatic/arginine selectivity filter (Ar/R SF) formed by four residues from TM2, TM5 and LE half-helix. As the name suggests, the Ar/R SF typically contains one arginine and at least one aromatic residue. The second constriction region is formed by the highly conserved Asn-Pro-Ala (NPA) motifs from LB and LE half-helices. These residues seem to play crucial role in the selectivity of the solutes that will be transported through the MIP channels. In the case of TbAQP2, there is deviation in both Ar/R SF and the NPA motif. Ar/R SF in TbAQP2 is formed by I-110, V-249, L-258 and L-264, all bulky hydrophobic residues. In structure-based generic numbering scheme, these residues correspond to the positions 2.49, 5.57, LE.47 and LE.53 respectively (1). In this numbering scheme, 2.49 indicates that this residue is in TM2 and it is one residue towards N-terminus from the most conserved residue in TM2 which is assigned the position 2.50. The conserved NPA motif in LB and LE half-helices are also substituted in TbAQP2. They are, respectively, NSA and NPS in LB and LE half-helices. In generic numbering, they correspond to LB.50 to LB.52 and LE.50 to LE.52, respectively. The conserved Pro is substituted by Ser in the first NPA motif present in LB and the Ala in second NPA motif is substituted by Ser. In one of the first studies, Uzcategui et al. showed that TbAQP2 transported glycerol and water when heterologously expressed in yeast and *Xenopus oocytes* (6). Alghamdi et al. carried out series of mutations in TbAQP2 and showed that specific mutations can influence pentamidine permeation (16). Molecular dynamics simulations reported in the same paper also showed that pentamidine permeation is energetically favorable in TbAQP2. With all residues in the Ar/R SF hydrophobic, the natural question is how could water be transported through the hydrophobic selectivity filter. How can the transport efficiency be compared with the prototype mammalian AQP1 channel? Can TbAQP2 be equally efficient in transporting glycerol? In this paper, the molecular mechanism of water and glycerol transport through the hydrophobic selectivity filter is investigated by employing molecular dynamics technique. Our results surprisingly show that TbAQP2 with hydrophobic selectivity filter can more efficiently transport water than that of AQP1.

## Materials and Methods

The structure of *Trypanosoma brucei* aquaporin 2 (TbAQP2) was predicted using AlphaFold (17). AlphaFold employs multiple sequence alignments (MSAs) and deep neural networks trained on structural databases to predict atomic-level protein structures. In our case, a template-based approach was used to model TbAQP2 where structurally similar aquaporins from the PDB (Protein Data Bank) (18) served as templates. The amino acid sequence of TbAQP2 retrieved from the UniProt database (UniProt ID: U5NJF5) (19)was used in AlphaFold to generate multiple models. The model with the highest-confidence structure based on predicted Local Distance Difference Test (pLDDT) scores was selected. The final tetrameric transmembrane aquaporin fold closely resembled its homologs validating the structure. The Alphafold-predicted model showed an RMSD of 0.66 Å with the recently determined experimental structure of TbAQP2 (PDB ID: 8JY7) (14). Energy minimization was further applied to resolve steric clashes and optimize local geometries.

### System set-up

We considered experimentally determined tetrameric structure of AQP1 (PDB ID: 1J4N) (20) and the modeled tetramer of TbAQP2. The systems were built using CHARMM-GUI Membrane Builder (21). The tetrameric structures were embedded in POPE lipid bilayers with 349 lipids (176 upper leaflet and 173 lower leaflet) (22). The systems were then solvated with TIP3P water molecules (23)ensuring complete hydration on both sides of the membrane. To maintain charge neutrality and physiological ionic strength, sodium (Na⁺) and chloride (Cl⁻) ions were added to the system. The final simulation box dimensions were 11.81344 × 11.81344 × 10.96007 nm^3^ ensuring that the protein-lipid complex was well separated from its periodic images and thus avoiding artificial interactions due to periodic boundary conditions (PBC).

### Simulation details

All simulations were performed using GROMACS-2018 (24). CHARMM36 force-field (25) was used for all systems. To begin with, energy minimization was performed in order to remove steric clashes, optimize atomic positions and ensure a stable starting conformation. Steepest descent minimization was conducted using the steepest descent integrator for up to 50,000 steps and the maximum force convergence criteria (emtol) was set to 1000 kJ/mol/nm. A Verlet cutoff scheme was applied for neighbor searching with a cutoff of 1.2 nm for both Coulombic (PME method) and van der Waals (VDW) interactions. After steepest descent minimization, a conjugate gradient minimization step was performed to further refine the energy landscape. The same convergence criteria as of steepest descent were used for this phase. Temperature of the systems was gradually increased from 50 K to 310 K in six steps over 1 ns. This was conducted under the canonical (NVT) ensemble ensuring that the system was thermally stabilized while maintaining a fixed volume. Temperature regulation was achieved using the Berendsen thermostat (26) which couples the system to an external heat bath adjusting atomic velocities to achieve the desired temperature. The pressure was controlled using the Berendsen barostat in a semi isotropic mode allowing pressure adjustments independently in the membrane plane (XY) and membrane normal (Z-axis) to preserve membrane properties like thickness.

Systems were first equilibrated under constant volume and temperature (NVT ensemble) for 24 ns ensuring that the protein, lipids and water molecules adjusted to thermal fluctuations while remaining confined within a fixed simulation box. Temperature control during this phase was achieved using the V-rescale thermostat (27) which maintains proper fluctuations for a true canonical ensemble. The system was divided into two heat-coupling groups that are SOLU_MEMB (protein-lipid complex) and SOLV (water and ions) each maintained at 310 K. Positional restraints were imposed on the protein backbone, side chains and lipid head groups which were gradually released in eight steps each ensuring smooth relaxation. First positional restraints of the lipid head groups were relaxed followed by the protein side chains and finally the protein backbone. Bond constraints on hydrogen atoms were maintained using the LINCS algorithm (28).

The next phase of equilibration consisted of 24 ns NPT simulation during which both temperature and pressure were regulated allowing for natural fluctuations in box dimensions to establish a stable lipid bilayer thickness and proper membrane density. Pressure coupling was achieved using the Parrinello-Rahman barostat (29) which accurately maintains pressure fluctuations and dynamically adjusts the simulation box dimensions. A semi-isotropic pressure coupling scheme was applied in which the membrane plane (XY) and membrane normal (Z) are controlled independently ensuring that the bilayer adapted to the embedded aquaporin without artificial distortions. The reference pressure was set to 1 bar. Temperature control remained consistent with the Nosé-Hoover thermostat (30) which maintains correct thermodynamic fluctuations by employing a chain of heat baths. Similar to NVT equilibration, positional restraints were imposed and gradually released over eight stages.

Once the system had attained thermal and pressure equilibrium, it was subjected to a 200 ns fully unrestrained run. No positional restraints were applied enabling the protein, lipid and solvent to interact freely and reach full dynamic equilibrium. Temperature control was maintained using the Nosé-Hoover thermostat (30), while pressure control continued using the semi-isotropic Parrinello-Rahman barostat (29) maintaining the membrane structure at 1 bar. By the end of this equilibration phase the system had achieved full relaxation. The stabilization of lipid bilayer properties (thickness, area per lipid & order parameter) and protein dynamics were examined to ensure that the systems were properly equilibrated. The production runs consisted of 1 microsecond (1 µs) simulation under NPT conditions. MD simulated structures were saved at every 10 ps for further analysis.

### Osmotic Water Permeability (*P_f_*) Calculation

By applying the collective diffusion model proposed by Zhu *et al*. (31), the osmotic permeability coefficient (*P_f_*) was determined for both systems by using the last 500 ns of the 1 µs trajectory to ensure accurate equilibrium conditions. The collective diffusion model defines the water transport process in terms of a collective coordinate *n(t)*, representing the total displacement of water molecules inside the channel over time. The time-dependent collective coordinate was calculated from the simulation trajectory using the equation.

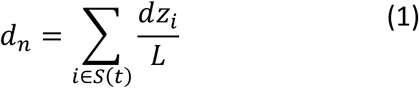

where L represents the length of the channel’s constriction region and *S*(*t*) is the set of water molecules inside the channel at time t. The quantity *dz_i_* corresponds to the one-dimensional displacement of each water molecule along the channel axis. The collective coordinate *n(t)* was computed at 10 ps intervals. For water molecules that entered or exited the channel during this period, only the portion of their displacement within the channel was included in the sum. To get the mean square displacement (MSD) of the collective coordinate, the full 500 ns trajectory was divided into 5000 independent sub-trajectories each having length 100 ps. The collective coordinate *n(t)* in each sub-trajectory was set to zero at t = 0 to remove offsets ensuring consistency across time windows. This was achieved by subtracting the initial value from the subsequent displacement values. The MSD of *n(t)* was then computed as:-

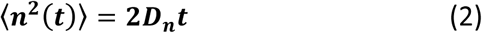

where *D_n_* represents the collective diffusion constant, calculated as the best-fit slope of the MSD curve. Using the computed collective diffusion constant (*Dn*), the osmotic permeability coefficient (*P_f_*) was obtained using the relation:-

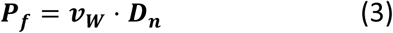

where ***v_W_*** is the molecular volume of a single water molecule taken as 3.0 × 10^−23^ cm^3^ (32).

### Calculation of potential of mean force (PMF) profile for water molecules

In order to compare the energetics of water transport between AQP1 and TbAQP2, the potential of mean force (PMF) profiles for water permeation were computed. The PMF profiles were generated for the last 500 ns of the production runs. A cylindrical sampling region was defined along the central pore axis of all four monomers to track the movement of water molecules through the channel. The cylinder was constructed with a length of 60 Å and a radius of 5 Å centered on the NPA motifs (z = 0 Å). The cylinder extended 30 Å towards the extracellular side and 30 Å towards the intracellular side covering the entire permeation pathway. To get the water occupancy along the pore, the cylinder was divided into 120 bins each with a thickness of 0.5 Å along the z-direction. The number of water molecules present in each bin was recorded across all frames of the trajectory (50,000 frames in total). The average water occupancy per bin was then calculated by dividing the total count in each bin by the number of frames providing a time-averaged representation of water distribution along the channel.

The PMF for a given bin z in an individual monomer was computed using the Boltzmann relation (33–34):

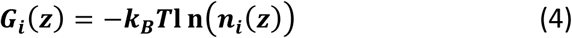

where ***k_B_*** is the Boltzmann constant (8.314 × 10⁻³ kJ/mol·K), T is the simulation temperature (310 K) and ***n***_***i***_**(*Z*)** is the average number of water molecules in bin z for monomer i.

Since the channel entrance and exit regions experience entropic effects that influence water distribution, a trapezoidal entropic correction was applied to the PMF values in these regions (35). The correction factor accounts for the difference in available cross-sectional area between the monomeric channel pore (*A*_*mono*_) and the cylindrical sampling region (*A*_*c*_) given by:-

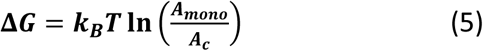

where *A*_*mono*_ is the cross-sectional area of a single monomer 923.781 Å² and *A*_*c*_ is the area of the cylindrical sampling region, which is 78.5 Å². Substituting these values the entropic correction was calculated as:

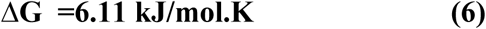

This correction was applied to the channel entrance and exit regions spanning the range −21 Å to +21 Å, corresponding to the distance between the lipid headgroups of the upper and lower bilayer leaflets (42 Å). The corrected PMF profiles for each monomer were computed individually and plotted to visualize the free energy landscape for each independent channel within the tetramer.

### Umbrella Sampling for Glycerol Transport

In order to determine the energetics of glycerol transport, umbrella sampling simulations (36)were conducted for water-transporting AQP1, glycerol-specific GlpF and TbAQP2. The mammalian aquaporin-1 (AQP1) and *Escherichia coli* aquaglyceroporin GlpF served as controls and used to compare the energetics of glycerol transport in TbAQP2. The CHARMM-GUI Membrane Builder suite was employed to construct membrane-embedded monomeric systems for each system. Each monomer was embedded in a POPE lipid bilayer ensuring a biophysically relevant membrane environment. The bilayer was fully solvated on both sides and counterions (Na+ and Cl−) were added to neutralize the systems. Energy minimization, simulated annealing and multi-stage equilibration were carried out following the standardized procedures as used for equilibrium simulations described above. The reaction coordinate for glycerol transport was defined as the distance along the z-axis (membrane normal) measuring the position of the glycerol molecule relative to the center of mass (COM) of the transmembrane region’s Cα atoms. To ensure a uniform initial starting position, the glycerol molecule was first placed 5 Å above the extracellular entrance of the pore and the surrounding water molecules within a 5 Å radius were removed using VMD to prevent steric clashes during the initial phase of substrate pulling. A steered molecular dynamics (SMD) simulation was conducted in GROMACS to gradually pull the glycerol molecule from the extracellular side to the intracellular exit. The pulling was performed along the z-axis using a constant velocity of 0.5 nm/ns with a force constant of 5000 kJ/mol/nm. This ensured a smooth and continuous transition of glycerol through the channel providing initial configurations for the umbrella sampling windows (37).

A total of 102 umbrella windows were defined along the reaction coordinate with each window spaced 0.04 nm apart. Within each window harmonic positional restraints were applied to keep glycerol fixed at its designated position along the reaction coordinate with a force constant of up to 2500 kJ/mol/nm. Each umbrella window was first equilibrated for 1 ns, allowing the system to relax around the restrained substrate. This was followed by a 5 ns production simulation during which umbrella sampling was conducted while maintaining harmonic restraints on the glycerol molecule. To restrict lateral diffusion, a flat-bottomed cylindrical potential was applied confining glycerol within a radius of 0.5 nm from the channel axis ensuring that transport remained along the physiologically relevant permeation pathway.

Potential of Mean Force (PMF) profiles were computed using the Weighted Histogram Analysis Method (WHAM) (38), implemented via the module in GROMACS. This method combines histograms from all umbrella windows, reconstructing the free energy landscape of glycerol transport through each system. To ensure optimal convergence, umbrella histograms were closely monitored. In regions where sampling was insufficient or convergence was slow, additional umbrella windows were introduced with higher force constants applied to improve statistical accuracy. To quantify statistical uncertainty, Bayesian bootstrapping (20 iterations) was employed providing a measure of confidence in the PMF results.

## Results

The aquaporin homolog TbAQP2 (UniProt ID: U5NJF5) shares 20.7% sequence identity (31.2% sequence similarity) with the prototype water channel AQP1 (UniProt ID: P47865) (Figure S1). As mentioned earlier, the experimentally determined TbAQP2 structure adopts the same aquaporin hour-glass helical fold although the sequence identity is very low. With unusual Ar/R SF with all four residues forming the Ar/R SF hydrophobic (Figure 1A and B), it is intriguing that TbAQP2 is also involved in water and glycerol transport as demonstrated by heterologous expression and functional studies (6). When we compared these two different aquaporin homologous structures within the transmembrane regions (PDB IDs: 1J4N vs 8JY7), the root mean square deviation (RMSD) is only 1.21 Å (Figure S2) and we considered only the Cα atoms for the purpose of calculating RMSD. We performed 1 μs equilibrium MD simulations on TbAQP2 and AQP1 systems as described in the Methods section to understand the molecular mechanism of water transport. We calculated the MD trajectories of RMSD, root mean square fluctuation (RMSF) and channel radius profiles for both TbAQP2 and AQP1 MD simulations. Our results show that the RMSD trajectories are stable for both TbAQP2 and AQP1 (Figure 2A). The average RMSD during the last 500 ns of production runs is 1.20 and 1.48 Å for TbAQP2 and AQP1 respectively indicating that two structures remained stable and close to the starting conformations. AQP1 deviated from the starting structure slightly more than that found for TbAQP2. We also analyzed RMSF plots for both simulations (Figure 2B). Apart from the large fluctuations in the N-terminus, both of them clearly show that the loop connecting the two halves of the hour-glass fold (TM1-TM3 and TM4-TM6) shows significant conformational variations in both cases. The other notable features in the RMSF profile are the large fluctuations in specific loops. The loop connecting TM1 and TM2 in AQP1 shows significantly larger fluctuation compared to the negligible fluctuation found for the same loop in TbAQP2. This is due to the fact that this loop is nearly ten residues longer in AQP1 compared to that in TbAQP2 (Figure S1, Figure 1C and 1D). The second loop showing differential dynamics is the loop connecting the half-helix LE and TM6. This is almost ten residues longer in TbAQP2 compared to that found in AQP1.

**Figure 1:**
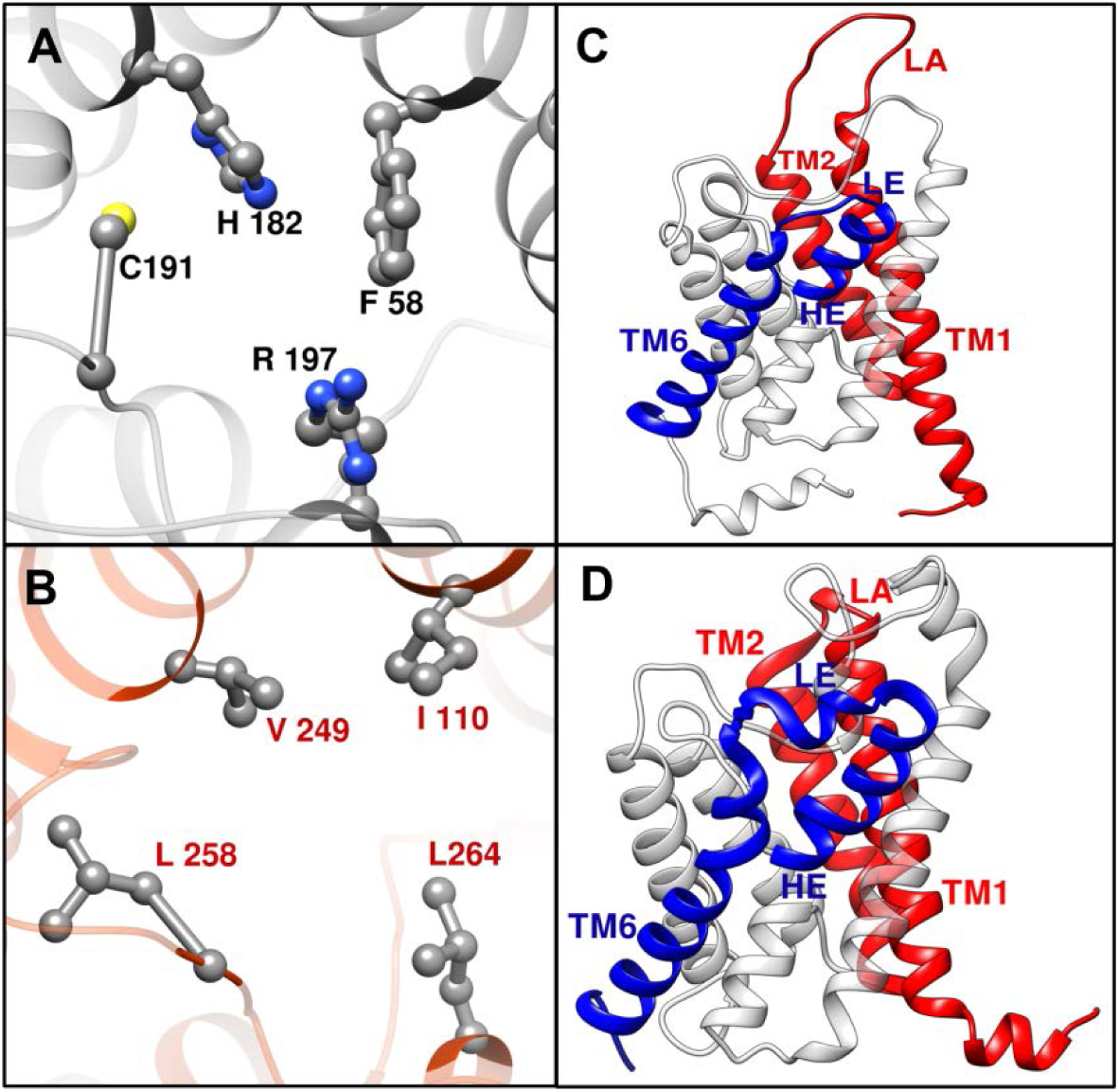
Residues forming the aromatic/arginine selectivity filter (Ar/R SF) for (A) AQP1 and (B) TbAQP2. The loop regions between (C) TM1 and TM2 and (D) LE and TM6 which are longer in one of the two aquaporin homologs are shown. The experimentally determined structures of AQP1 (PDB ID: 1J4N) and TbAQP2 (PDB ID: 8JY7) have been plotted in which the segments of TM1-TM2 and LE-TM6 are shown in red and blue colors respectively.

**Figure 2:**
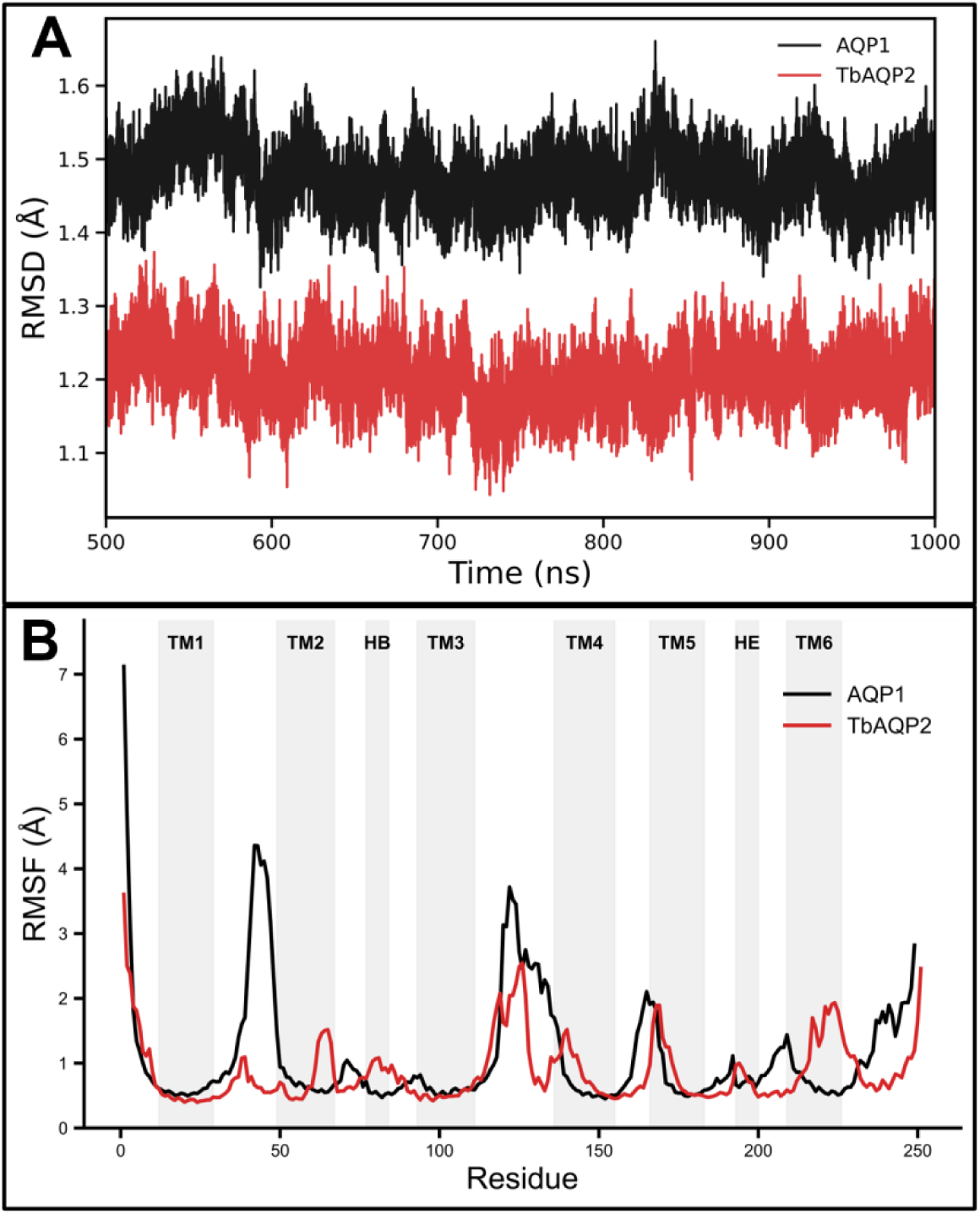
(A) MD trajectories of RMSD for AQP1 and TbAQP2 systems as a function of time. Each point is the average calculated for all the four monomers. (B) RMSF analyses for AQP1 and TbAQP2. The transmembrane regions and half-helices are indicated by gray bars.

### Comparison of channel characteristics

Since the transport of solutes depends upon both the size and chemical nature of channel constriction, we calculated the channel radius profiles of TbAQP2 and AQP1 using the program HOLE (Figure S3) (39). The experimentally determined crystal structures were used for this purpose. Channel radius profiles show that the channel is wider in both Ar/R SF and the NPA regions in TbAQP2. In Ar/R SF region, the channel radius is larger by more than 0.6 Å in TbAQP2 while in the NPA region, the radius is greater by 0.4 Å. We also calculated the temporal radius profiles for both TbAQP2 and AQP1 (Figure 3). Temporal radius profiles will give an idea how the channel width varies at a specific point as the simulation progressed. Although the TbAQP2 channel radius is relatively bigger than that found for AQP1 in the narrowest regions, the intriguing question is how the ar/R SF with all hydrophobic residues can allow water molecules to pass through. To answer this question, we have also plotted the average channel radius profiles with standard deviation calculated for the last 500 ns of production runs (Figure 4). Comparison of temporal radius profiles of TbAQP2 and AQP1 shows that the AQP1 channel appears to be narrow for a much longer region during the simulation period compared to TbAQP2. Average channel radius profiles exhibit a clear widening of TbAQP2 channel spanning a region which covers both constriction regions, namely Ar/R SF and NPA motifs. The average channel radius is ∼1.9 Å in the Ar/R SF region of TbAQP2 while it is 1.4 Å for AQP1.

**Figure 3:**
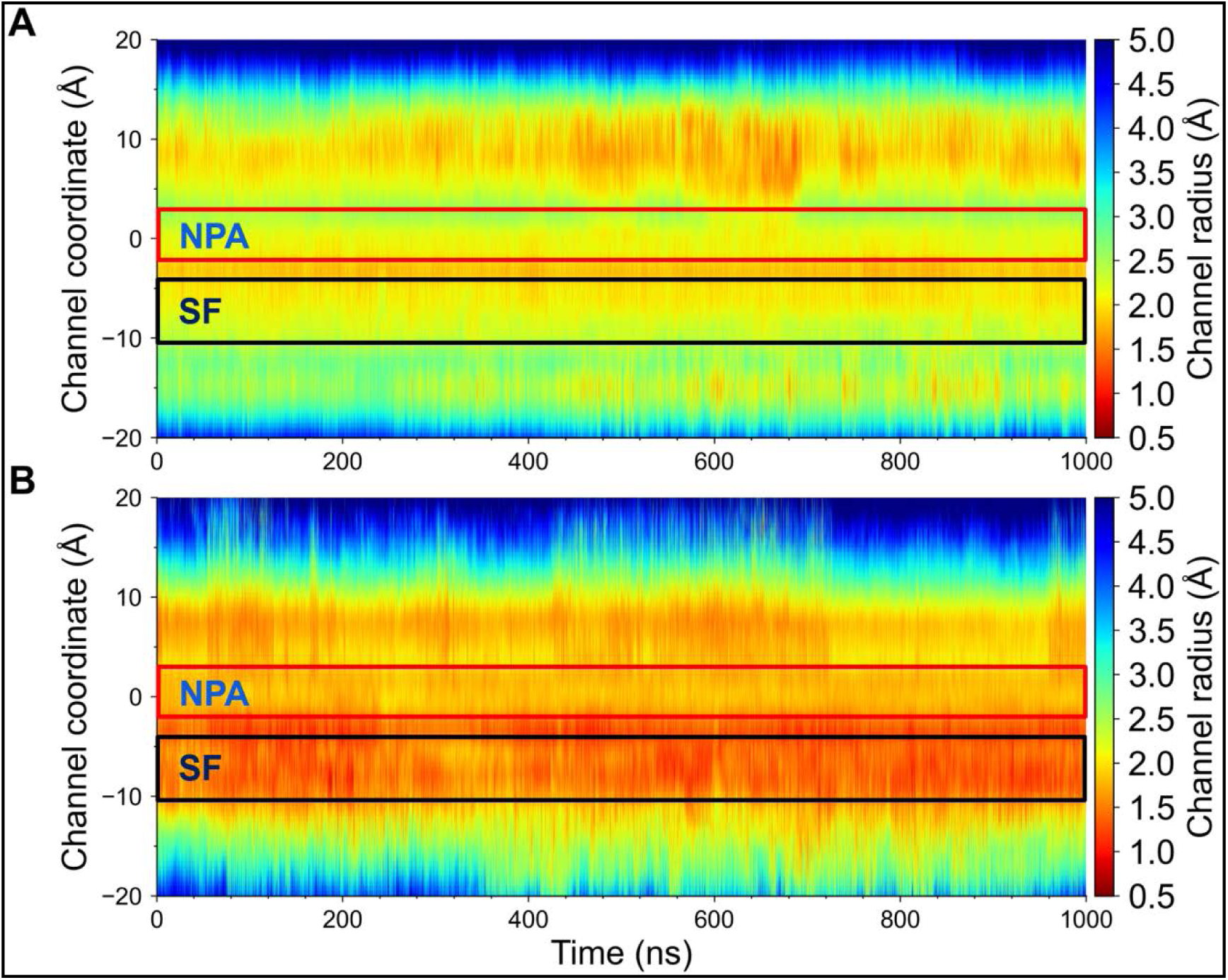
Temporal channel radius profiles for (A) TbAQP2 and (B) AQP1. Each point in the trajectory is the average calculated across four monomers. The Ar/R SF region and the NPA motif regions are indicated in the figure.

**Figure 4:**
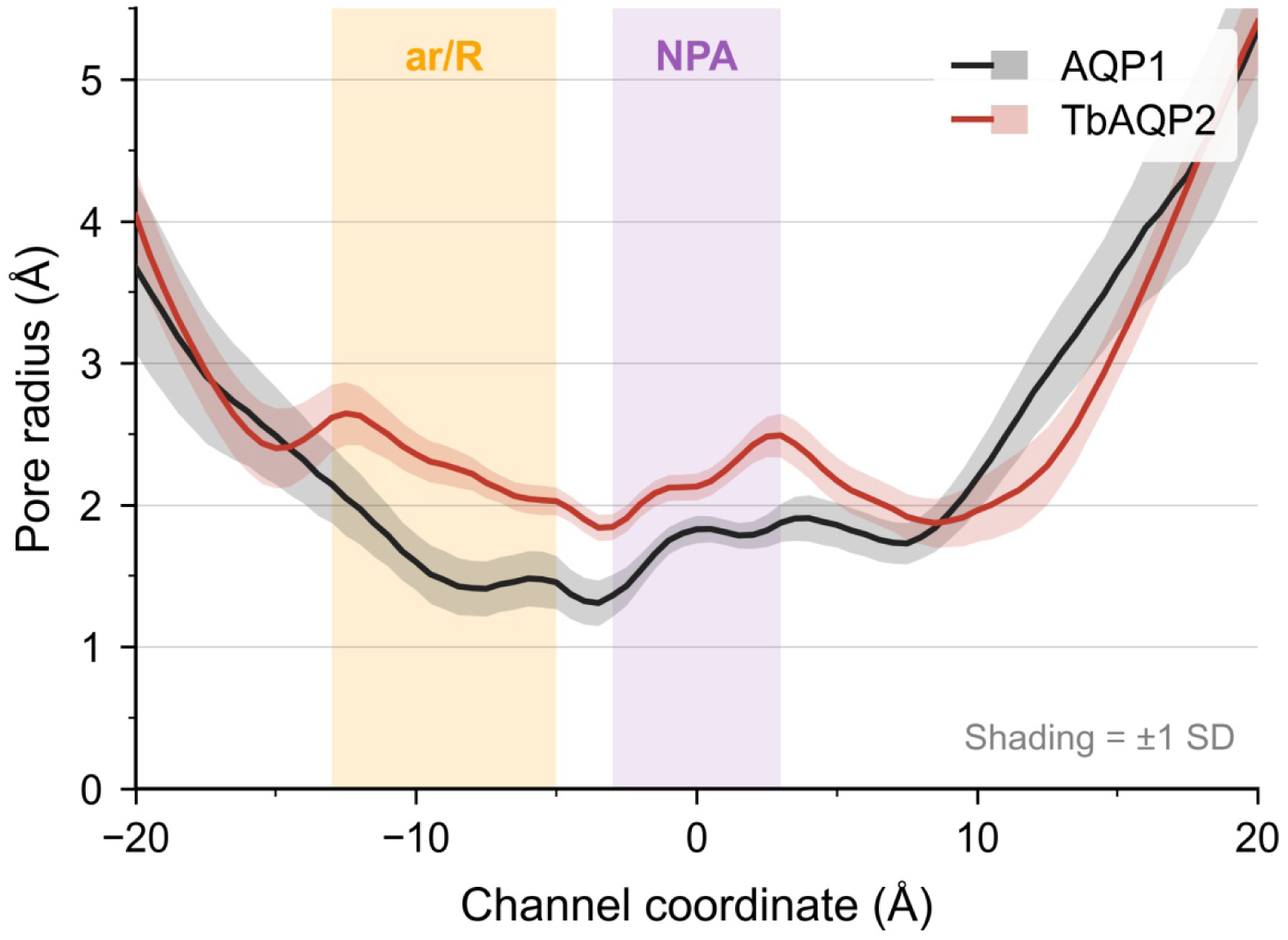
Average channel radius profiles calculated during the last 500 ns production runs for AQP1 and TbAQP2. The Ar/R SF region and the NPA motif regions are indicated by orange and purple bands respectively.

### Water transport properties: TbAQP2 vs AQP1

We then monitored the total number of water permeation events for the last 500 ns of the production runs for both TbAQP2 and AQP1. Since aquaporins facilitate bidirectional water transport, water molecules could traverse the channel in both directions. A permeation event was defined if a water molecule fully crosses the pore from one side to the other. To track permeation, a cylindrical region with a radius of 5 Å was defined along the pore axis for each monomer. This cylinder extends 15 Å above the NPA motif and 5 Å below it, covering the functional pore region. The cylinder was further divided into bins of 1 Å thickness along the z-axis allowing precise tracking of water movement in each bin. A water molecule was considered to have permeated the channel if it entered the cylindrical region from one side, traversed all the bins and exited on the opposite side. Water molecules that appeared on the opposite side due to periodic boundary conditions (PBC) effects were discarded to avoid artificial counting of permeation events.

Number of water permeation events for each of the four monomers during the last 500 ns of production run is summarized for both TbAQP2 and AQP1 in Table 1. Data related to residence time inside the channel for the water molecules are also provided in the same table. Residence time provides an idea of the duration that a water molecule spends within a specific region of the pore while traversing from one side to the other. This analysis gives insights into the interaction strength of water molecules within the channel’s constriction sites particularly at the Ar/R SF and NPA motif regions.

**Table 1:**
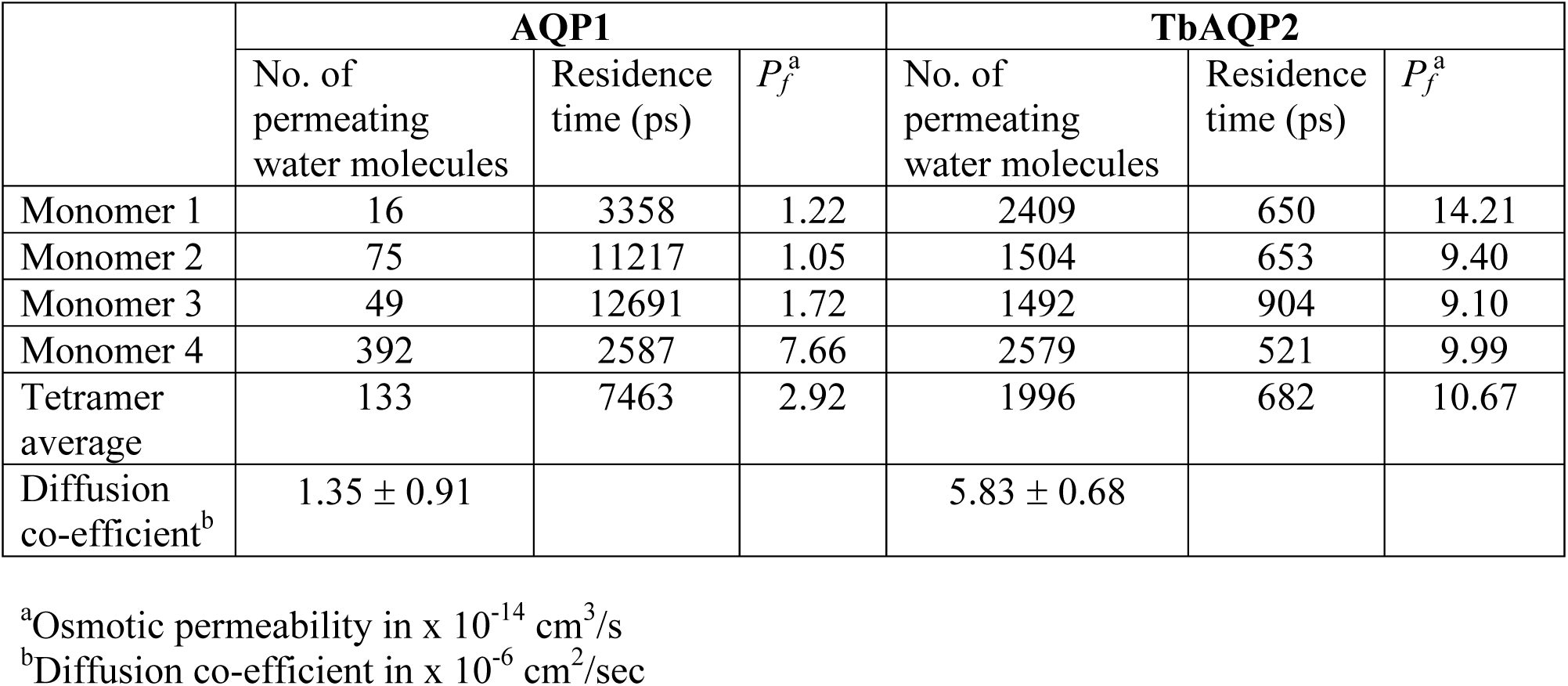
Comparison of water transport properties between TbAQP2 and AQP1 for the last 500 ns of production runs.

While the average number of water molecules transported through all four monomers is close to 2000 in TbAQP2, it is only 133 for AQP1. Each monomer in TbAQP2 permeates close to 1500 or more water molecules (Table 1). However, the maximum number of 392 water molecules transported in AQP1 is by monomer #4 and monomer #1 permeates smallest number of 16 water molecules. Thus, the number of water molecules transported by TbAQP2 is more than one to two orders of magnitude higher than that found for AQP1. This is also confirmed by calculating osmotic water permeability (*p_f_*) for each monomer in TbAQP2 and AQP1 as described in the Methodology section. The <*p_f_*> is only 2.92 × 10^−14^ cm^3^/s across all four monomers in AQP1 while it is 10.67 × 10^−14^ cm^3^/s in TbAQP2, clearly indicating that more water molecules are transported in TbAQP2 compared to AQP1 (Table 1 and Figure 5A).

**Figure 5:**
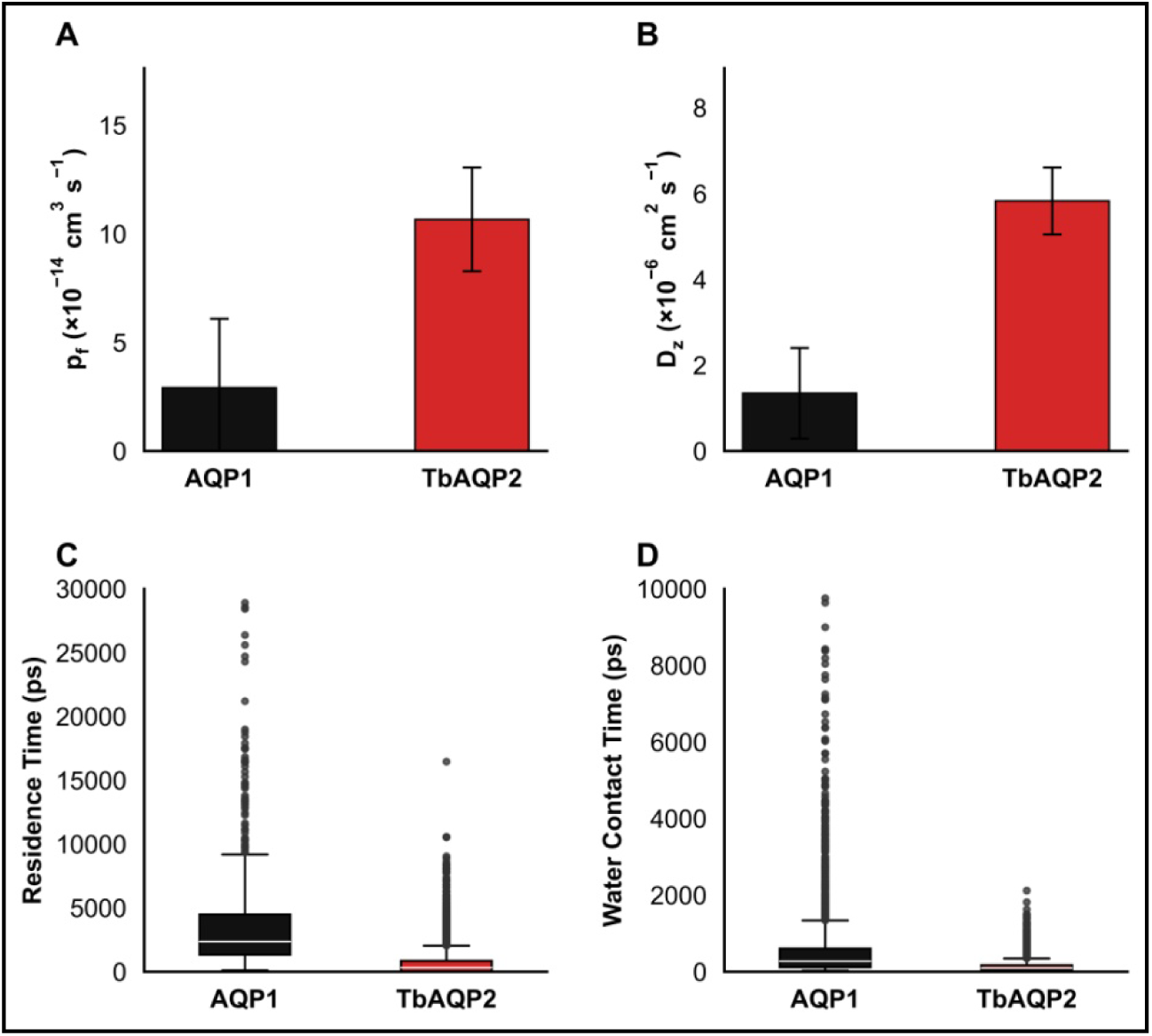
Comparison of water transport properties between AQP1 and TbAQP2. Box plots showing average (A) osmotic water permeability, (B) diffusion coefficient, (C) residence time of permeating water molecules and (D) water contact time with the selectivity filter residues. Averages were calculated for the last 500 ns of production runs.

We also calculated the residence time of water molecules inside the channel between the two narrow constriction regions. The residence time within the two constriction regions was calculated for each permeating water molecule that successfully crossed the channel in the last 500 ns of the simulation. The region we considered was defined as the span between the Ar/R SF and the NPA motif, covering the narrowest and most functionally critical segment of the pore. The residence time for each water molecule was obtained by subtracting the entry time from the exit time within the Ar/R SF-NPA regions. To ensure accuracy, abrupt fluctuations caused by PBC effects were filtered out and only gradual and physically meaningful permeation through the pore were considered. While average residence time for AQP1 is close to 7500 ps, it is only 682 ps for water molecules in TbAQP2 (Table 1 and Figure 5C). The smaller residence time indicates that the water molecules move more than 10 times faster in TbAQP2 than that found for AQP1. This could be due to two possible factors. Our channel radius profile analysis clearly indicates the region between Ar/R SF and NPA motifs in TbAQP2 is broader compared to AQP1 allowing the faster movement of water molecules. The second factor is that there are no specific functional groups in Ar/R SF region that are involved in stronger interactions within TbAQP2 with the permeating water molecules. Such interactions if present, would have slowed down the movement of water molecules. To further substantiate this point, we calculated the diffusion coefficient of permeating water molecules within the two channels. The diffusion co-efficient of water in TbAQP2 varies between ∼5.0 to ∼6.6 × 10^−6^ cm^2^/s in all four monomers while in AQP1, it varies from ∼0.6 to ∼2.8 × 10^−6^ cm^2^/s indicating that water molecules in TbAQP2 moves two to five times faster than that found for AQP1 (Table 1 and Figure 5B).

### PMF profiles of water: TbAQP2 vs AQP1

We calculated the PMF profiles for the permeating water molecules for both TbAQP2 and AQP1 (Figure 6). TbAQP2 consistently showed lower energy throughout the channel for the water molecules to pass through. AQP1 displayed energy barriers before and after the NPA motifs and the barrier is maximum between Ar/R SF and NPA regions. The free energy difference in the energy barrier between AQP1 and TbAQP2 is about 2 to 3 kJ/mol indicating that water molecules are more easily transported through TbAQP2 channel compared to AQP1. We also looked at the average number of water molecules in the two constriction regions. In general, higher number of water molecules is found in Ar/R SF compared to NPA motif region (Figure S4). Between the two systems, average number of water molecules is higher in TbAQP2 compared to AQP1 in both the constriction regions.

**Figure 6:**
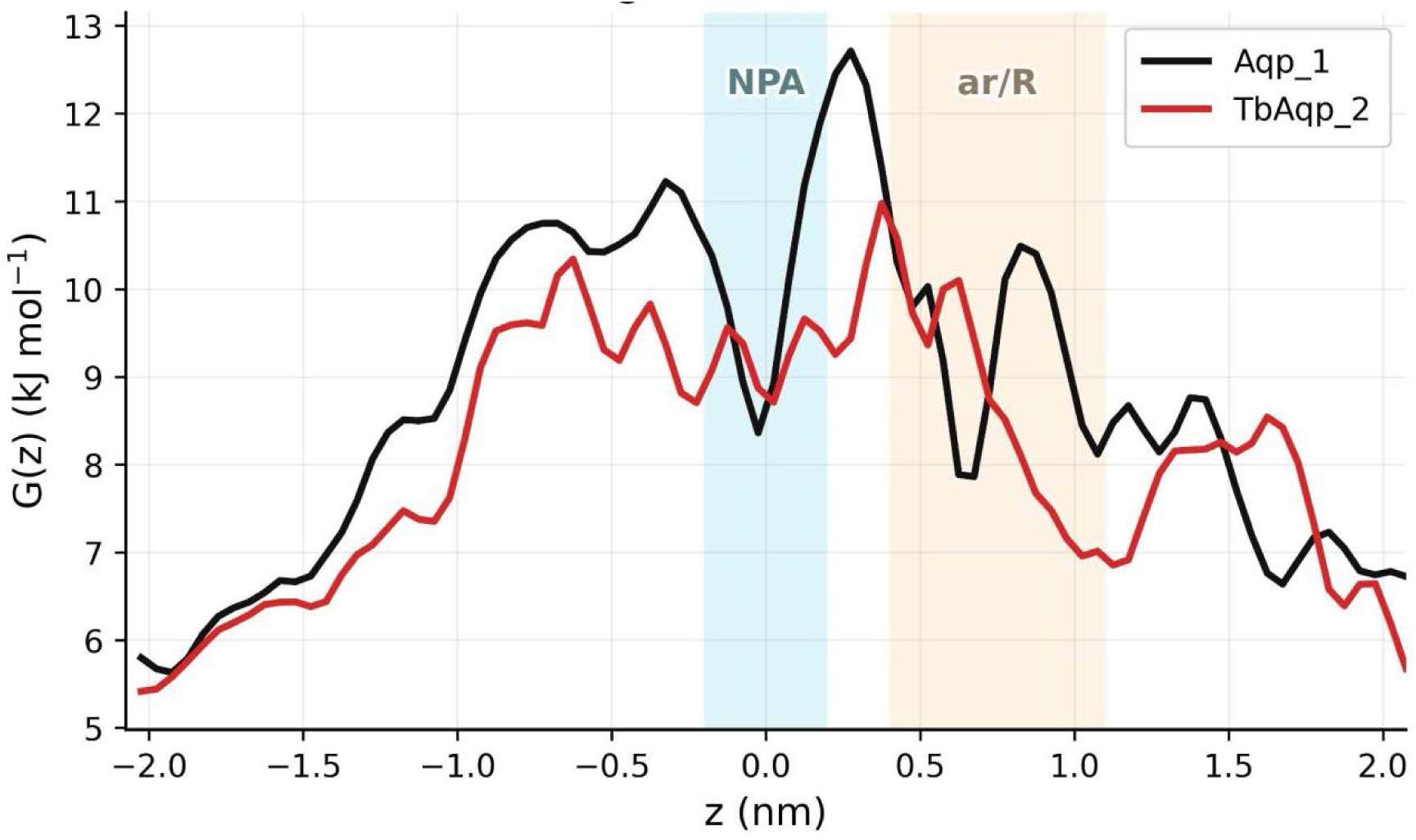
Potential of Mean Force (PMF) profiles of water molecules for AQP1 and TbAQP2. The NPA motif and the Ar/R SF regions are shown as blue and orange bands respectively

We further investigated the factors that are responsible for the large number of water molecules transported through TbAQP2. As mentioned earlier, one reason could be the larger pore size at the Ar/R SF region. The other reason could be the Ar/R SF residues in AQP1 could favorably interact with water molecules and these interactions could be responsible for the water molecules to be retained for a longer time. On the other hand, all Ar/R SF residues are hydrophobic in TbAQP2 and hence no specific interactions could take place with water molecules. We monitored the water molecules that come in contact with the Ar/R SF residues in both AQP1 and TbAQP2. We also examined the contact time of water molecules that are within 4 Å from one of the Ar/R SF residues. An average number of ∼13 water molecules are in contact with the Ar/R SF residues (F58, H182, C191 and R197) of AQP1 when all four monomers were considered. A subset of these water molecules permeates the channel and their average interaction time with Ar/R SF residues varies between 580 to 1200 ps in AQP1 (Figure 5D). In the case of TbAQP2, an average number of 16.5 water molecules are in contact with at least one of the four hydrophobic residues of Ar/R SF which is higher than that found for AQP1. Among them, the average contact time of permeating water molecules with Ar/R SF residues of TbAQP2 is about 100 to 180 ps across all monomers. This interaction time range is much lower than that of AQP1. The reason for longer contact time in AQP1 can be explained by finding out if these water molecules are interacting with the Ar/R SF residues especially with H182 and R197. We found out that average number of 4.25 water molecules is involved in hydrogen bond interactions with His and/or Arg functional groups. We used the geometric criteria involving distance between donor and acceptor (*d*(D…A) < 3.5 Å) and the angle between Donor-Hydrogen….Acceptor (*θ*(D-H…A) > 90°). This is about 25% of all water molecules that are in contact with Ar/R SF residues. Additionally, we also found out that significant number of all water molecules that are in contact with Ar/R SF residues in AQP1 and TbAQP2 make contact with more than one Ar/R SF residues. This analysis could explain why the water molecules are stuck at the Ar/R SF region in AQP1. Due to the hydrophobic side-chains in Ar/R SF region in TbAQP2, there are no interactions between permeating water molecules and the Ar/R SF residues and hence the residence time of permeating water molecules is very low. In the case of AQP1, water molecules could interact with Ar/R SF residues in the narrow constriction region, make hydrogen bonds that could slow down their permeation. As a result, large number of water molecules could permeate TbAQP2 compared to AQP1. A snapshot of water molecules in contact with Ar/R SF residues in AQP1 and TbAQP2 is shown in Figure 7.

**Figure 7:**
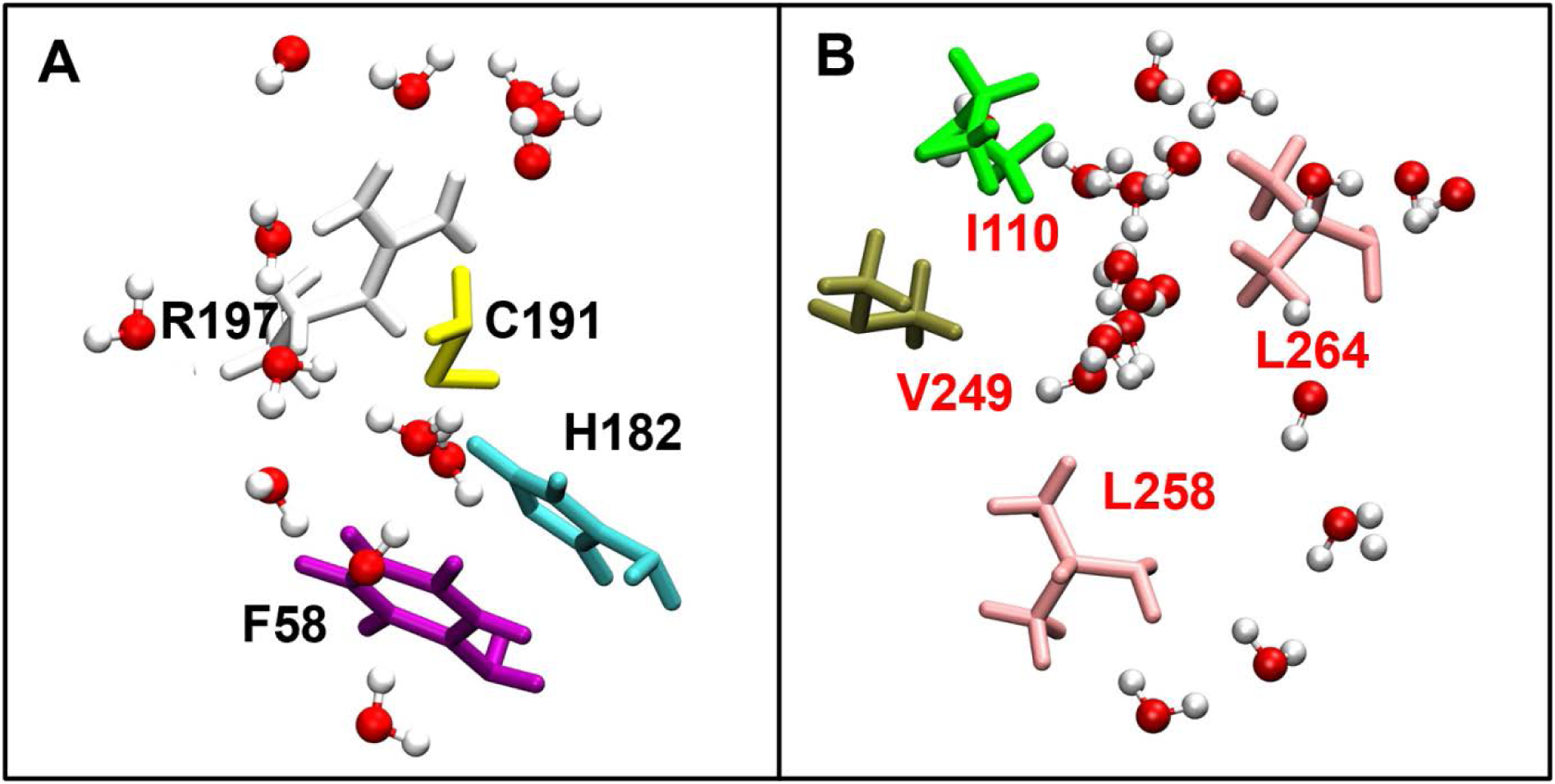
MD snapshot of Ar/R SF residues of (A) AQP1 and (B) TbAQP2 along with water molecules that are within 4 Å from one of the selectvity filter residues.

### Glycerol permeation through TbAQP2

We used umbrella sampling to compute the potential of mean force (PMF) profile for TbAQP2 as described in the Methods section. We followed the same protocol to calculate the PMF profiles of glycerol for AQP1 and GlpF as AQP1 and GlpF as prototype examples of water-transporting and glycerol permeating MIP homologs, respectively. Comparison of all three PMF profiles (Figure 8) clearly shows that AQP1 has huge energy barriers, one near Ar/R SF region and a much bigger one near the NPA region. GlpF shows the most favorable PMF profile for glycerol permeation. The PMF profile for TbAQP2 is comparable to GlpF but shows slightly higher energy barriers compared to that of GlpF consistently, especially near Ar/R SF region and NPA motif regions. The most favorable interaction for glycerol in TbAQP2 occurs in the NPA region in which the glycerol oxygens are involved in hydrogen bond interactions with backbone C=O group of His-128 and side-chain carboxyl group of Glu-77 (Figure 9). We then investigated how the Ar/R SF residues distinctly interact with the permeating glycerol molecule in TbAQP2 and GlpF. An interesting observation was that the glycerol molecule changes its orientation in the Ar/R SF region in TbAQP2 while it remains same in GlpF (Figure 10). When we examined the contacts made by permeating glycerol with the Ar/R SF residues, glycerol participated in electrostatic interactions with SF residues Arg and Trp, and made hydrophobic contacts with Phe and Trp in GlpF. In the case of TbAQP2, glycerol had to tumble and make hydrogen bond with the backbone C=O group of Ala-259, while the other hydrophobic residues of Ar/R SF region enabled hydrophobic interactions providing suitable environment for the glycerol to pass through. Thus the Ar/R SF residues are completely of different nature in GlpF and TbAQP2, they manage to provide compensating interactions that help to provide the glycerol an environment suitable to cross the narrow constriction.

**Figure 8:**
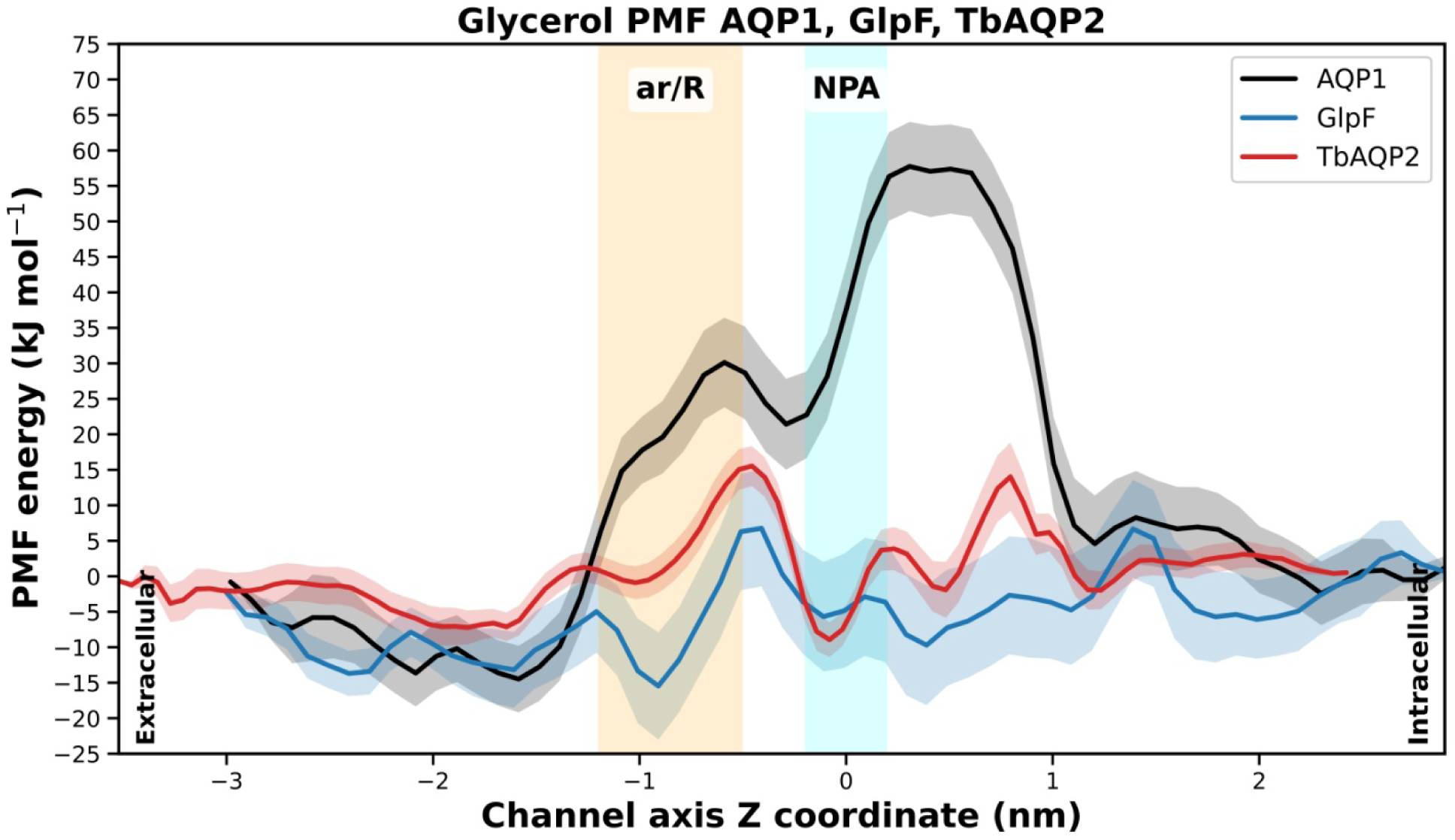
Potential of mean force (PMF) profiles of glycerol permeating through AQP1, GlpF and TbAQP2

**Figure 9:**
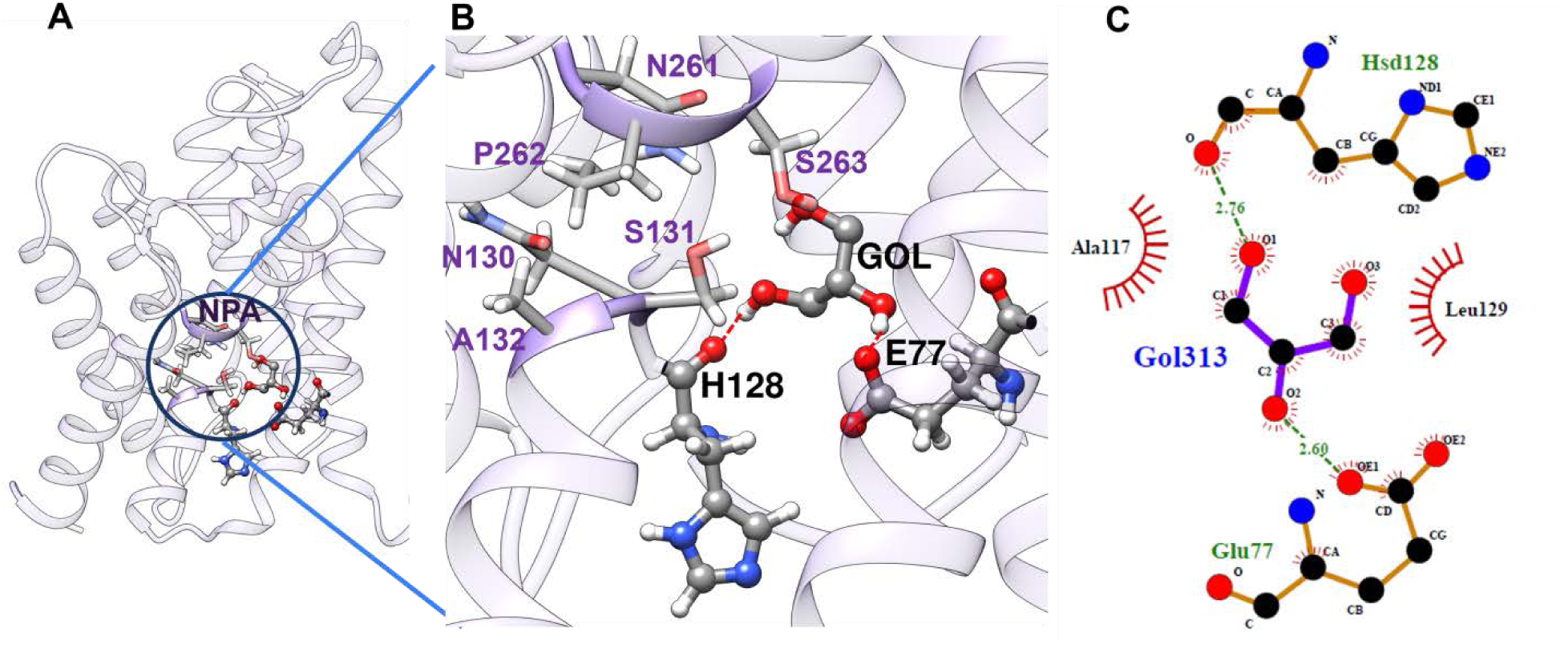
(A and B) The most favorable interactions between Glycerol and the NPA region. Glycerol forms hydrogen bonds with the backbone carbonyl group of His-128 and the side-chain carboxyl group of Glu-77. (C) LigPloT representation of Glycerol’s interactions with residues in the vicinity of NPA region of TbAQP2.

**Figure 10:**
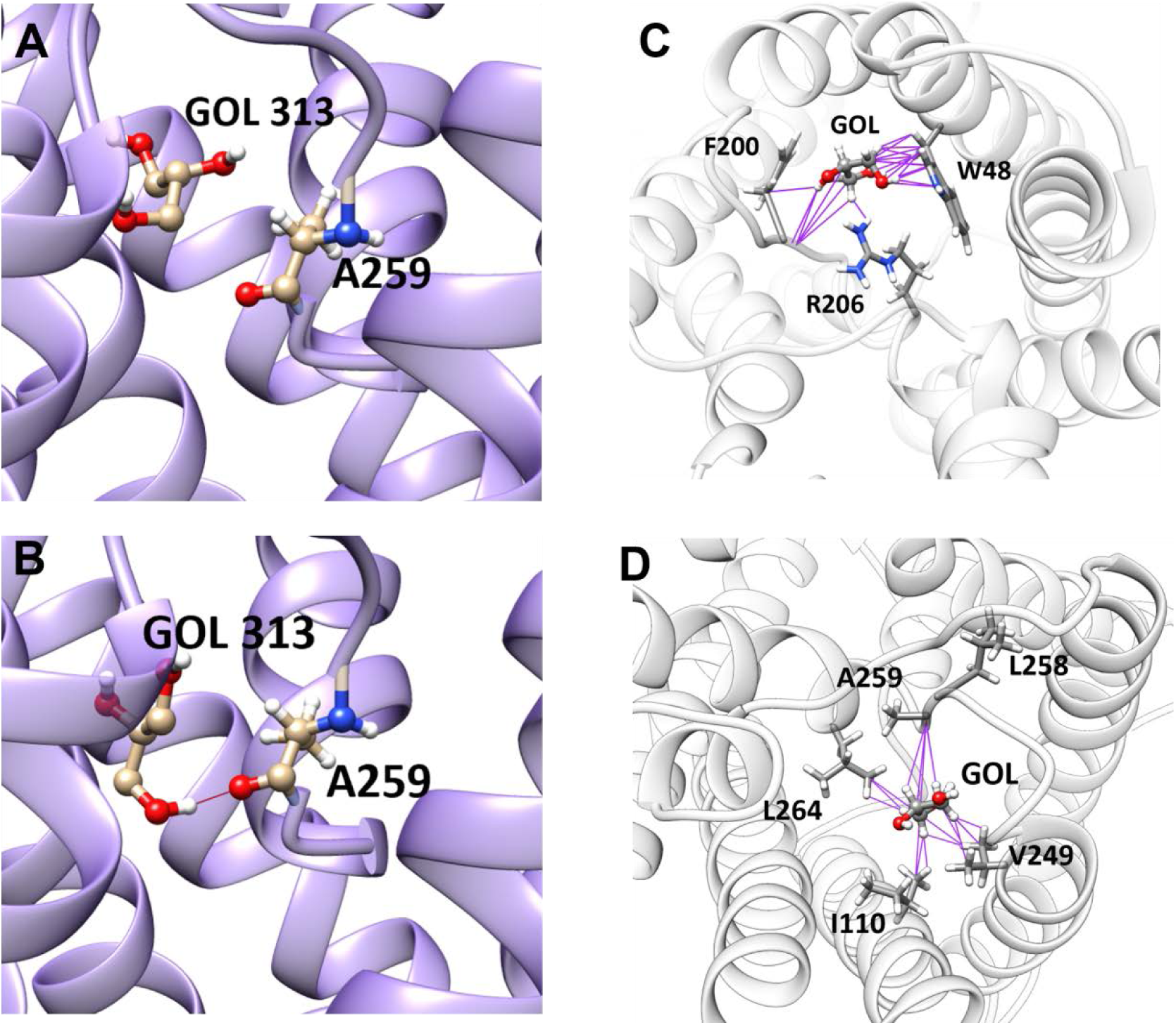
Interactions of glycerol molecule with the Ar/R SF residues in TbAQP2. (A and B) Glycerol molecule changes its orientation and interacts with the backbone carbonly oxygen of A259 in TbAQP2. Glycerol molecule’s interactions with the Ar/R SF residues of (C) GlpF and (D) TbAQP2. Ar/R SF residues which are within 4 Å of Ar/R SF residues are connected by purple lines. Both polar and non-polar contacts are made in both systems but with different residues.

## Discussion

Pathogenic parasitic aquaporins are least exploited as drug targets and are waiting to be explored for developing anti-parasitic drug compounds specific for particular diseases. Three-dimensional structures of all the MIP homologs determined to date have the same hourglass helical fold. The functional importance of aromatic/arginine selectivity filter and the NPA motifs in selecting the solutes to be transported have been studied using mutation experiments of MIP homologs from different species. For example, replacement of His-180 and Arg-195 in the Ar/R SF of rat AQP1 by Ala or Val enlarged the pore radius and resulted in glycerol transport (40). Single or double mutants resulting in removal of the positive charge residing in the arginine of Ar/R SF allow proton transfer indicating the vital role of Arg residue in the selectivity filter in preventing proton transport. Another study used single amino acid substitutions in Ar/R SF of AQP1, AQP3 and AQP4 and concluded that substrate selectivity depends on several factors including pore size, polarity and the nature of solute (41). Several studies investigated the role of Ar/R SF in different subfamilies of plant MIPs (42–45). Computational studies suggested that the arginine in Ar/R SF helps to inhibit the proton transport (46). Recently, Chen et al. used solid state NMR and computational techniques to investigate the importance of Arg in the Ar/R SF and concluded that Arg is indispensable to maintain the electrostatic distribution and to establish single file water chain for efficient water transport (47). The above studies focused on the role of Arg in Ar/R SF in the selectivity of the solutes. In the present study, the TbAQP2 lacks not only Arg but also aromatic residues. By having all four hydrophobic residues in the Ar/R SF, the outstanding question is how is it possible for TbAQP2 to transport water. Hydrophobic residues are supposed to repel water molecules and one would have obviously assumed that TbAQP2 will transport other solutes but not water. However, the experimental studies clearly established that TbAQP2 is involved in the transport of water, glycerol and other solutes. How do we explain water transport through hydrophobic SF and what is the molecular mechanism are the main goals of this study. In addition to water transport, we also investigated the molecular mechanism of glycerol transport. Our studies show that two important factors played vital role in the transport properties. TbAQP2 channel is as wide as the prototype glycerol channel GlpF. Our equilibrium MD simulation studies surprisingly resulted in a larger water transport than the typical AQP1 channel. The number of waters transported through the channel and the residence time of the transported waters indicated that almost one order of higher water molecules are transported through TbAQP2 and the residence time is much lower in TbAQP2 in comparison with AQP1. Due to the hydrophobic nature of Ar/R SF residues and the wider constriction, the water molecules that enter the channel immediately leave resulting in higher number of water molecules that are transported faster. In other words, TbAQP2 seems to be more efficient water channel than AQP1. This mechanism is reminiscent of transport observed in synthetic hydrophobic nanopores and carbon nanotubes, where reduced friction and weak wall-water interactions lead to anomalously fast water transport despite the absence of favorable interactions (48–50). This is understandable as the parasite has to encounter different osmotic environment in the host and the quick transport of water in either direction will help the parasite to survive. As far as the glycerol transport is concerned, our umbrella sampling studies showed that the PMF profiles of TbAQP2 and GlpF are almost similar. A closer examination reveals that as the glycerol enters the channel, its interactions with the channel residues differ from GlpF. This can be explained as the sequence similarity between these two channel is very less (33% sequence identity and 49% sequence similarity). Our study explains the molecular mechanism of water and glycerol transport in TbAQP2 aquaporin channel with hydrophobic selectivity filter. This knowledge can be exploted in the development of any channel blockers that can block the transport of water and/or glycerol.

## Data Code and Availability

All molecular dynamics simulation input files, including system configurations, topology files, and parameter files are available in a public repository at https://github.com/ramasubbu-sankar/TbAQP2-MD-Project. Additional data are available upon reasonable request.

## Acknowledgements

This work is supported by funding from Anusandhan National Research Foundation (ANRF) of Government of India (Sanction Order No. CRG/2023/002313). We gratefully acknowledge the availability of PARAM Sangathan and High Performance Computing facility at IIT-Kanpur. PMP thanks IIT-Kanpur for a Fellowship. We thank all our lab members for the discussion.

## Author Contributions

PMP performed simulations and data analysis. RS conceived and designed the research. RS wrote the manuscript and all authors approved the final version of the manuscript.

## Supporting Material

Supporting material is available online

