## Supplementary Figures for "Molecular mechanism of water and glycerol transport through hydrophobic selectivity filter in the aquaporin homolog of *Trypanosoma brucei*"

### SUPPORTING INFORMATION

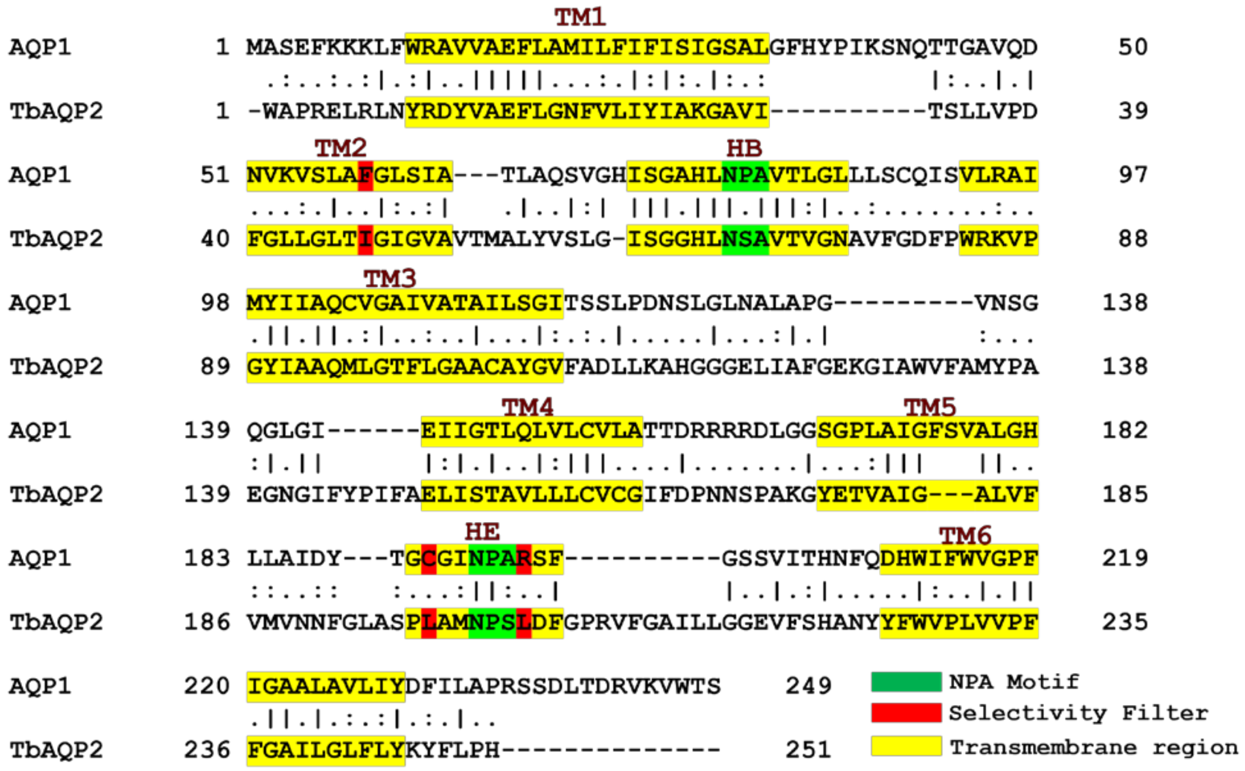


**Figure S1:** Pairwise sequence alignment of AQP1 (UniProt ID: P47865)) and TbAQP2 (UniProt ID: U5NJF5) sequences. Transmembrane regions, NPA motifs and the residues forming the Ar/R SF region are highlighted in colors.

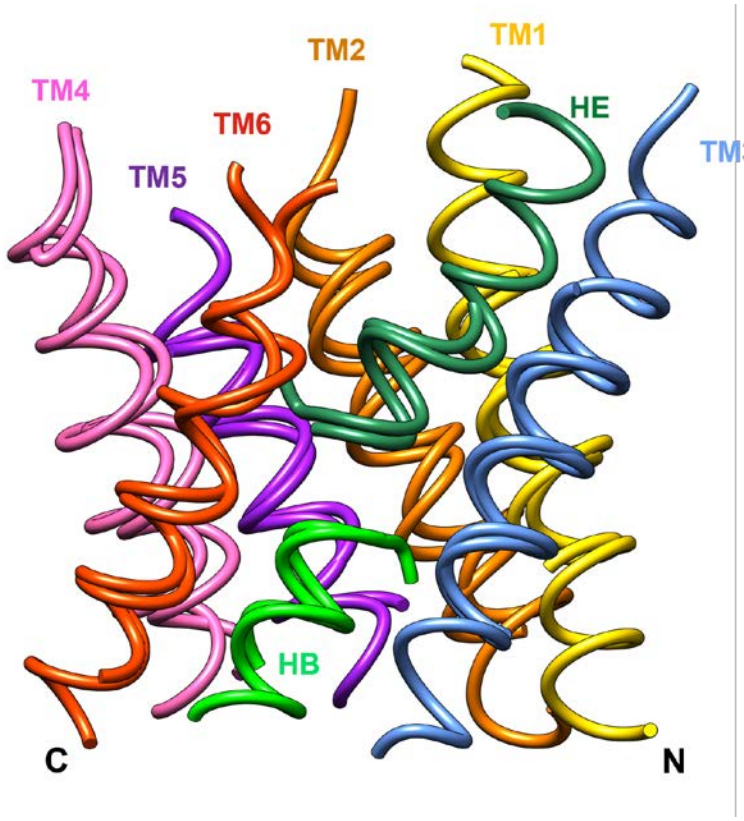


**Figure S2**: Superposition of transmembrane helical regions of AQP1 (PDB ID: 1J4N) and TbAQP2 (PDB ID: 8JY7). Each TM helix is shown in one color.

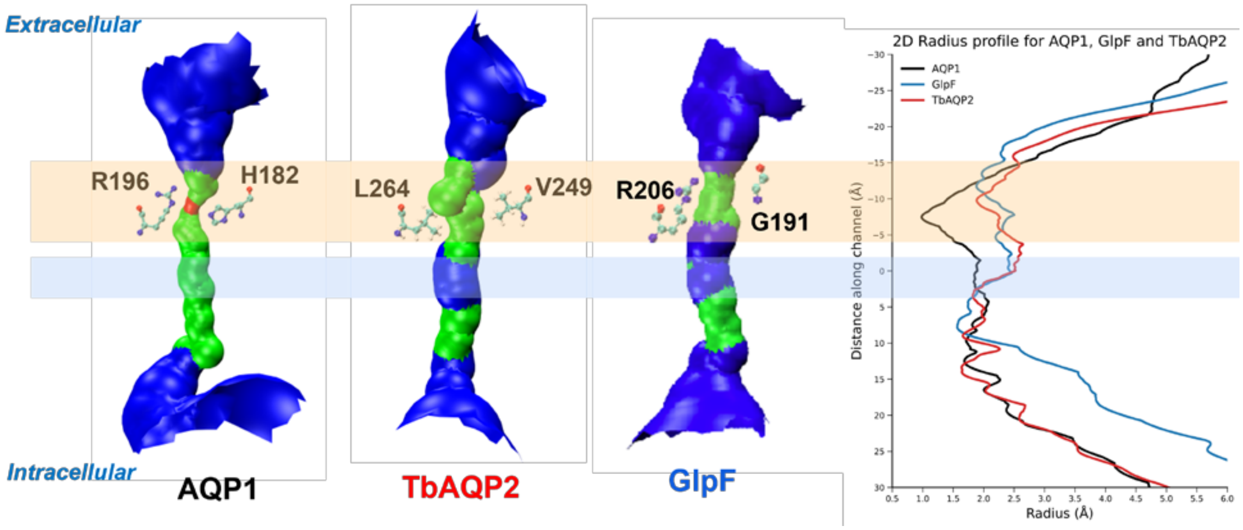


**Figure S3**: Channel radius profiles generated using the program HOLE for the three aquaporin homologs. The aromatic/argine selectivity filter region and the NPA motif regions are shown in orange and blue bands respectively. The arginine and aromatic residues in AQP1 and the residues at the equivalent positions in TbAQP2 and GlpF are shown in stick representation. The 2D channel radius profiles of all three channels are shown at the right.

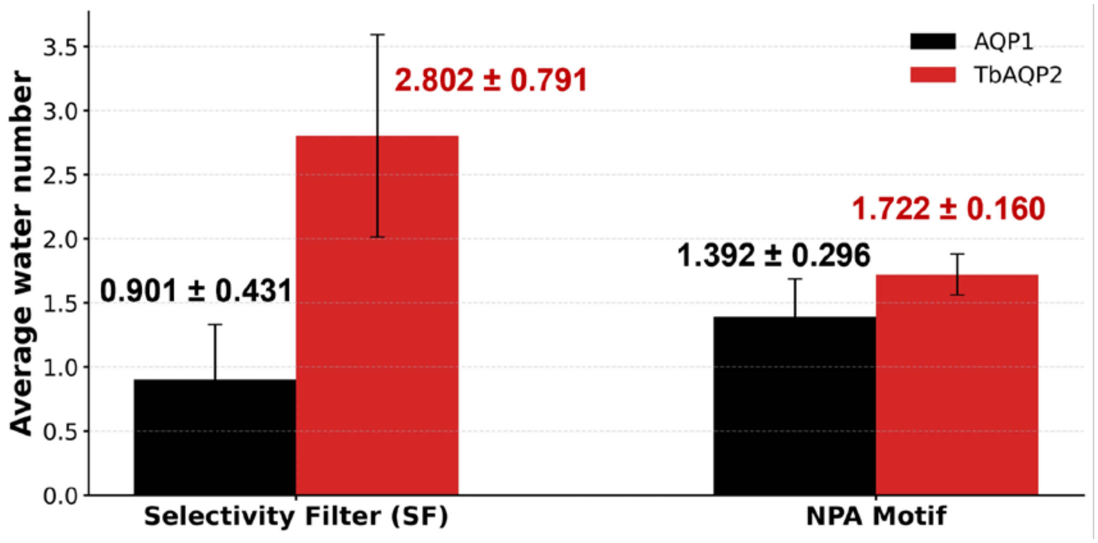


**Figure S4**: Average number of water molecules in the aromatic/arginine selectivity filter region and NPA motif regions shown for AQP1 and TbAQP2.
